# Elucidating the Contribution of OVLT Glutamatergic Neurons to Mineralocorticoid Hypertension in TASK^−/−^ Mice

**DOI:** 10.1101/2025.06.11.659218

**Authors:** Luo Shi, Kun Liu, Ke Zhao, Wei He, Yongqiang Chen, Xue Zhao, Yanxuan Wei, Jinting Chen, Sheng Wang, Fang Yuan

## Abstract

**Background:** Aldosterone overactivity intensifies central sodium sensitivity and sympathetic output, driving salt-sensitive hypertension, but specific mechanisms remain incompletely defined. Herein, we aimed to explore the role of organum vasculosum of the lamina terminalis glutamatergic neurons (OVLT^Glut^) and their hyperexcitability mechanisms in hyperaldosteronism-associated hypertension.

**Methods:** Adult age matched male TASK^−/−^ mice (primary aldosteronism model) and wild-type controls (TASK^+/+^) mice were used. Neuronal excitability was assessed via patch-clamp techniques. Arterial blood pressure (BP) monitored via telemetry or carotid catheterization. Chronic drug delivery used minipumps. RNA-seq/qPCR profiled gene expression, and intracerebroventricular hypertonic saline tested sodium sensitivity.

**Results:** In TASK^−/−^ mice, heightened OVLT^Glut^ activity increased sympathetic outflow and hypertension, mitigated by OVLT^Glut^ neuron ablation. Optogenetic activation of these neurons or their paraventricular nucleus (PVN) / rostral ventrolateral medulla (RVLM) projections acutely elevated BP, with ablation reducing BP selectively in TASK^−/−^ mice. Aldosterone dependence of OVLT^Glut^-PVN/RVLM neuron hyperactivity was evident in both TASK^−/−^ mice and TASK^+/+^ mice with chronic aldosterone infusion. Aldosterone chronic infusion enhanced central sodium pressor effects, that were nullified by OVLT^Glut^-PVN/RVLM neuron lesioning. RNA-seq indicated that aldosterone-induced ion channel expression spectrum changes, including potassium channels and the epithelial sodium channel, underlie the neuronal hyperexcitability.

**Conclusion:** Overactivation of OVLT^Glut^ neurons contributes to hypertension in TASK^−/−^ mice through regulation of OVLT^Glut^-PVN/RVLM circuits. The hyperexcitability of these neurons, possibly due to aldosterone-induced changes in ion channel expression spectrum, contribute to hypertension by amplifying central sodium sensitivity.

## Introduction

Hypertension remains a global health emergency, contributing to over 10 million annual deaths and ranking as the leading modifiable risk factor for cardiovascular morbidity and mortality worldwide^1^. Salt-sensitive hypertension (SSH) represents a distinct phenotype affecting approximately 50% of essential patients, characterized by aberrant renal sodium handling and a rise in arterial BP with higher dietary salt consumption^1,2^. Despite decades of investigation, the precise mechanisms linking dietary sodium excess to sustained BP elevation remain incompletely elucidated^2,3^. Sympathetic hyperactivity is a critical contributor to SSH, a complex disorder where excessive dietary sodium intake leads to disproportionately elevated BP^4^. Evidence from both salt-sensitive humans^5,6^ and animal models^7^ shows that a high-salt diet only mildly increases plasma or cerebrospinal fluid NaCl levels (2 to 6 mM). If NaCl is indeed a contributor to tonic sympathoexcitation, a question is prompted: how can such slight increments in NaCl lead to considerable and continuous activation of the sympathetic nervous system? Other factors likely enhance sympathoexcitation from elevated plasma NaCl levels^8^. Meanwhile, osmoreceptors or Na⁺ receptors do not significantly reset with prolonged hyperosmolality, allowing sustained neural activation^9,10^. Recent investigations have identified AngII and aldosterone as critical neurohumoral amplifiers that establish a permissive milieu for NaCl-mediated sympathetic activation^8,11^. Landmark studies by Alan Kim Johnson^2,4,12^, John W. Funder^13^, Sean D. Stocker^14,15^, Joel C. Geerling^16,17^, and Arthur D. Loewy^18,19^ have demonstrated that the interaction between elevated AngII or aldosterone and high salt intake induces neuroplastic changes in the central nervous system, amplifying hypertension and sympathetic nervous system activity, a pivotal process in SSH development. Across all these studies, further clarification of SSH is essential, as its mechanisms remain incompletely understood and current animal models fail to accurately replicate human phenotypes.

Central nervous system integration of osmotic and volume signals occurs primarily in the lamina terminalis, a forebrain region comprising the median preoptic nucleus (MnPO), subfornical organ (SFO), and OVLT^20^. These structures, particularly the SFO and OVLT, lack a complete blood-brain barrier, enabling direct detection of endocrine, sodium and osmolytes from the blood and cerebrospinal fluid. Neurons within the OVLT and SFO are inherently sensitive to precise fluctuations in osmolality or extracellular NaCl^21–23^. Importantly, human brain imaging studies utilizing PET or fMRI reveal that acute hypertonic saline infusion, raising plasma osmolality by about 1% to 2%, triggers activation of the lamina terminalis^24,25^. Lesion studies show that OVLT is involved in multiple hypertension models, including Grollman renal hypertensive models^26^, and Goldblatt 2K1C hypertensive rats^27^. In line with this concept, acute optogenetic or chronic chemogenetic activation of OVLT neurons elevates sympathetic nerve activity and arterial BP^28^. Anatomical tracing shows direct projections from OVLT/SFO neurons to the paraventricular nucleus (PVN)^21^. Recent studies indicate that the PVN integrates osmotic signals from the OVLT into vasopressinergic signals^29,30^ or relays them to the RVLM via glutamatergic transmission, enhancing sympathetic activity^31,32^. Although limited data exist regarding the activity of OVLT in salt-sensitive models (or humans), using in vivo single-unit recordings, Stocker’s group revealed that the neuronal discharge of NaCl-responsive OVLT neurons was elevated in Dahl-Salt Sensitive rats fed a high-salt diet^33^.

Despite these advances, critical questions remain: (1) What are the anatomical substrates and molecular mechanisms underlying aldosterone’s modulation of salt-induced sympathoexcitation? (2) How does chronic aldosterone excess alter neuroplastic within these circuits? (3) Which neurotransmitter systems mediate sodium balance-to-sympathetic activity coupling? Given the high prevalence and major health hazards of SSH, it is imperative to seek answers to these questions through established and innovative technical approaches. As a frequent cause of secondary hypertension, primary aldosteronism is involved in the pathogenesis of salt sensitivity hypertension^13,34^. Short-term sodium restriction can more effectively lowers BP in PA patients than in those with essential hypertension, indicating heightened salt sensitivity in the former condition^35,36^. The TASK^−/−^ mouse model^37^, which recapitulates key features of PA including hyperaldosteronism, hypernatremia, hypokalemia, and hypertension, provides a unique platform to investigate these questions. By focusing on the OVLT glutamate system (OVLT^Glut^) in this model, we aim to dissect the molecular mechanisms of hypertensive response sensitization and neural circuitry through which aldosterone excess translates sodium retention into sustained sympathetic outflow.

## Material and methods

### Experimental animals

The TASK^−/−^ mice, a well-defined mouse model of primary aldosteronism generated by the global knockout of TASK-1 and TASK-3 channels as previously described^37,38^, were generously provided by Dr. Douglas Bayliss from the University of Virginia. VGlut2-Cre mice (stock number 016963) were purchased from The Jackson Laboratory. C57BL/6 mice (TASK^+/+^) were obtained from Vital River Laboratory Animal Technology *Co. Ltd.* (Beijing, China). All mice used in this study were age-matched males (12 to 15 weeks old, 23 to 30 grams) due to sex differences in the TASK^−/−^ phenotype^37^. TASK^−/−^ mice and VGlut2-Cre mice were separately backcrossed onto a C57BL/6 background for at least 10 generations. Mice were maintained on a 12-hour light/dark cycle in a temperature- and humidity-controlled vivarium (22 ± 2 °C; 50% humidity) with free access to food and water in a pathogen-free animal care facility at Hebei Medical University. All animal studies were performed after randomization. Animal use followed the NIH Guide for the Care and Use of Laboratory Animals and adhered to the Guide for the Care and Use of Laboratory Animals of the Chinese Association for Laboratory Animal Science. It was approved by the Animal Care and Ethics Committee of Hebei Medical University (IACUC-Hebmu-P2023366).

### Anesthesia and analgesia

For all surgical procedures, anesthesia was induced using 100% oxygen/2-3% isoflurane delivered at 1.5 L/min using an Animal Anesthesia System (RWD Life Science, China), and followed by maintenance at 100% oxygen /1.5% isoflurane via an inhalation mask. During these surgeries/procedures, the level of anesthesia was monitored by checking the eye blink reflex and a reaction to hind paw pinch, and was adjusted if necessary. For recovery surgery, ketorolac (4 mg/kg, i.p., RS37619, MCE, China) was administered immediately prior to the surgical procedure and also during the post-surgical recovery period (every 8 h for 3 days). Antibiotics (ampicillin, 125mg/kg, i.p., HY-B0522, MCE, China) were administered once daily for 3 days post-surgery. For euthanasia, mice were administered a lethal dose of urethane (4 g/kg, i.p., HY-B1207, MCE, China), followed either by decapitation or by transcardial perfusion with saline and 4% paraformaldehyde (PFA, Shenggong Biotech, *China*).

### BP Recording

The method for recording arterial BP in conscious and anesthetized mice is referenced from our previously published paper^38^. Briefly, mice were anesthetized (isoflurane inhalation) and telemetric ECG transmitters (HD-X11, Data Sciences International) were implanted subcutaneously. Mice recovered for at least five days before continuous 24-hour monitoring with hourly measurements at 1 kHz. For anesthetized mice, a PE50 tube was inserted into the left common carotid artery, connected to a pressure transducer. Data was recorded at 0.1 kHz and analyzed with LabChart software (PowerLab26T, AD Instruments).

### Stereotactic Adeno-Associated Virus Injections

The detail method refers to our previous published paper^39^. Mice were anesthetized with isoflurane and secured in a stereotaxic frame. Virus injections were performed using a micropipette attached to a syringe pump, with a rate of 60 nL/min and held for 5 minutes. For OVLT neuron ablation, AAV9-CaMKIIa-EYFP-taCasp3-ETVp was used, while controls received AAV9-CaMKIIa-EYFP (100 nL). For optogenetic activation, AAV9-hSyn-DIO-ChR2-EYFP or AAV9-hSyn-DIO-EYFP was injected into the OVLT of VGlut2-Cre mice (100 nL). To ablate OVLT^Glut^-PVN/RVLM neurons, AAVretro-CaMKIIa-GFP-2A-Cre was injected into the PVN or RVLM (80 nL per side), with AAV9-CAG-DIO-taCasp3-ETVp-WPRE-pA or AAV9-CAG-DIO-WPRE-pA (control) injected into the OVLT (100 nL). After experiments, animals were euthanized and histologically examined for virus expression. All viruses (titer > 1 × 10¹²/mL) were purchased from Brain Case Biotechnology Co., Ltd. (China) and stored at –80 °C.

### Immunohistochemistry and RNAscope

The detail method refers to our previous published paper^39^. Mouse brains were processed, sectioned at 25 µm, blocked, and incubated with primary and secondary antibodies. Sections were imaged after mounting. Antibody sources are in Table S1. For RNA in situ hybridization, the RNAscope Multiplex Fluorescent Reagent Kit V2 (ACDBio) was used with the RNAscope probe *Slc17a6-C2* (catalog# 319171-C2). Brain sections (13 µm) were treated with hydrogen peroxide, rinsed in PBS, incubated with primary antibodies overnight, and followed by protease treatment and hybridization as per the standard RNAscope protocol. After secondary antibody incubation, slices were washed and mounted with DAPI-Fluoromount-G (Southern Biotech).

### Photostimulation

The detail method refers to our previous published paper^39^. For in vivo photostimulation of ChR2-expressing neurons, a 200 µm optical fiber connected to a 473 nm LED light source (Newdoon Inc., China) was used. The light power output at the fiber tip was calibrated to 10 mW for all experiments, measured with a power meter. A fiber optic cable and cannula were attached to the stereotaxic arm to precisely target and stimulate the OVLT, PVN, and RVLM. Blue light (473 nm) was delivered to these regions under the specified parameters to observe BP changes. BP was recorded in the anesthetized state as described above.

### Whole-cell patch clamp recordings

The whole-cell patch clamp recordings were performed using the standard patch clamp technique as previously described^40,41^. Briefly, whole-cell patch clamp recordings were performed on 250 µm brain slices containing the OVLT. Slices were prepared in a cold sucrose solution and incubated in artificial cerebrospinal fluid (aCSF) at 34°C for 30 minutes, then at room temperature for 1 hour. Recordings were conducted at room temperature with a 2 mL/min aCSF perfusion. Neurons were visualized and recorded using a fluorescence microscope and Axon Multiclamp 700B amplifier. Data was acquired at 10 kHz with borosilicate glass electrodes (3–6 MΩ) filled with an intracellular solution. Resting membrane potential (RMP) was recorded post-whole-cell configuration, with compensation for capacitance and series resistance. Neurons with unstable or depolarized potentials below - 40 mV were excluded. Electrophysiological data analysis was performed with pClamp10. Action potential (AP) parameters were measured following 500 ms current injections (−15 pA to 55 pA, 5 pA steps). Input resistance was measured by the voltage response to a −15 pA hyperpolarizing current injection. AP parameters included threshold (dV/dt = 10 mV/ms), amplitude, half-width, afterhyperpolarization (AHP) amplitude, and AHP time. CNQX (catalog# 0190, Tocris Bioscience) at 10 mM was perfused with aCSF to validate the neurotransmitter type discharged by ChR2-expressing OVLT^Glut^ neurons.

### Chronic Aldosterone Treatment and RNA-Seq Analysis

Alzet minipumps (1007D) were soaked in saline overnight before implantation in isoflurane-anesthetized mice. An incision was made between the scapulae, and the minipumps filled with either vehicle (5% EtOH) or aldosterone (HY-113313, MCE, China, 900 μg/mL) was placed subcutaneously^17^. This approach can maintain the aldosterone concentration in the blood at approximately 1000 pg/mL, which is close to the aldosterone level in TASK^−/−^ mice^17,38^. The incision was closed with sutures, and an antibiotic ointment was applied. Mice were housed individually, monitored daily for 7 days, then anesthetized and decapitated for OVLT tissue collection. Total RNA was extracted using Trizol Reagent and then sent to Gene Denovo Biotechnology (Guangzhou, China) for further sequencing. The detail method refers to our previous published paper^40^. Bioinformatic analyses were conducted using the Omicsmart platform (https://www.omicsmart.com/). The raw data are available on Mendeley Data (https://data.mendeley.com/datasets/typp4cw8bz/1). Single-cell RNA sequencing dataset of the OVLT in wild-type mouse was acquired from the GEO database (GSE154048, http://www.ncbi.nlm.nih.gov/geo/).

### Intracerebroventricular infusion

To perform direct intracerebroventricular infusions of hypertonic solution, a 27-G guide cannula (RWD Life Science, China) was stereotaxically positioned at the lateral ventricle (AP, −0.5 mm; ML, 1.0 mm; DV, −2.5 mm; relative to the bregma) and was fixed to the skull. Hypertonic solution (450 mM [Na^+^]) was prepared by adding an appropriate volume of NaCl to aCSF. Intracerebroventricular infusions were performed using 33-G internal cannulas connected to a syringe pump (Pump 11 Elite, Harvard Apparatus) for 10 minutes at a rate of 0.4 μL/min. The Na^+^ concentration and infusion rate were derived from the literature^42^.

### RT-qPCR

OVLT tissue was dissected from brain slices. RNA was extracted with Trizol and reverse-transcribed to cDNA using HiScript Q-RT Mix (R223-01, Vazyme, China). qPCR was conducted with ChamQ SYBR Master Mix (Q711-02, Vazyme, *China*), normalized to GAPDH. Primers are detailed in Table S2.

### Statistics

Statistical analysis of data was performed using GraphPad Prism v9 and the results were expressed as the means ± standard error or median with interquartile range, as appropriate. Difference between the two groups was analyzed by Student’s *t* test with normal distribution or by Mann-Whitney U test with skewed distribution. The differences among multiple groups were analyzed using one-way ANOVA followed by Tukey’s post hoc test or by two-way ANOVA with Bonferroni post hoc test. For the comparison of 24-hour dynamic BP and HR, Fisher’s LSD post hoc test was employed. P values less than 0.05 were considered statistically significant.

## Results

### Ablation of OVLT^Glut^ Neurons Reduces Hypertension and Sympathetic Nerve Activity in TASK^−/−^ Mice

Previous research indicates that TASK^−/−^ mice develop hypernatremia and hypertension, which can be mitigated by a low-sodium diet^43^. The OVLT plays a crucial role in regulating BP by sensing extracellular Na^+^ concentrations, pointing to it as a potential key target of hypertension in TASK^−/−^ mice. To test our hypothesis, we compared cFos immunoreactivity in the OVLT between TASK^−/−^ and TASK^+/+^ mice. A significant increase in cFos-positive cells in TASK^−/−^ mice versus controls (Figs.1A-C) indicates heightened activation level of OVLT neurons in this model. Given the known predominance of glutamatergic neurons (∼80%) in this brain region^44^, we employed RNA fluorescent in situ hybridization to determine neuronal phenotype. Strikingly, 92.5±3.1% of cFos^+^ neurons co-expressed the vesicular glutamate transporter 2 gene (*Slc17a6*), indicating that the heightened OVLT activity in TASK^−/−^ mice primarily originates from glutamatergic neurons (OVLT^Glut^, Figs.1 D, E). These findings establish hyperactivation of OVLT^Glut^ as a potential target contributing to hypertension in this model. Activation of OVLT is known to increase water intake, so we observed whether TASK^−/−^ mice would drink more water. Unexpectedly, we found no significant difference in the total water consumption over 24 hours between the two genotypes (Fig.S1). A plausible explanation is that chronic OVLT activation in TASK^−/−^ mice could induce neural network compensatory mechanisms, thus avoiding sustained thirst.

To further determine the role of OVLT^Glut^ neurons to hypertension in TASK^−/−^ mice, we selectively ablated these neurons using AAV-CaMKII-Cre-EGFP and AAV-CAG-DIO-taCasp3-TEVp co-injection, inducing apoptosis via Cre-mediated caspase-3 expression, as reported previously^45^. Mice received AAV-CaMKII-Cre-EGFP alone as control (Fig.1F). Four weeks post-AAV injection, immunostaining showed abundant EGFP-positive neurons in OVLT of control mice, whereas taCasp3-ablated mice showed nearly no EGFP-positive neurons, confirming effective ablation (Figs.1G, H). Continuous radio telemetry monitoring demonstrated that OVLT^Glut^ ablation did not affect mean arterial pressure (MAP) or its circadian rhythm in TASK^+/+^ mice (Figs.1I, J). In contrast, TASK^−/−^ mice exhibited significant MAP reduction post-ablation (by approximately 6 mmHg), though values remained elevated compared to wild-type controls (Figs.1I, J). Notably, heart rate (HR) remained unchanged in both genotypes following neuroablation (Figs.1K, L). These results imply that OVLT^Glut^ neurons are crucial for sustaining hypertension in TASK^−/−^ mice.

**Figure 1.**
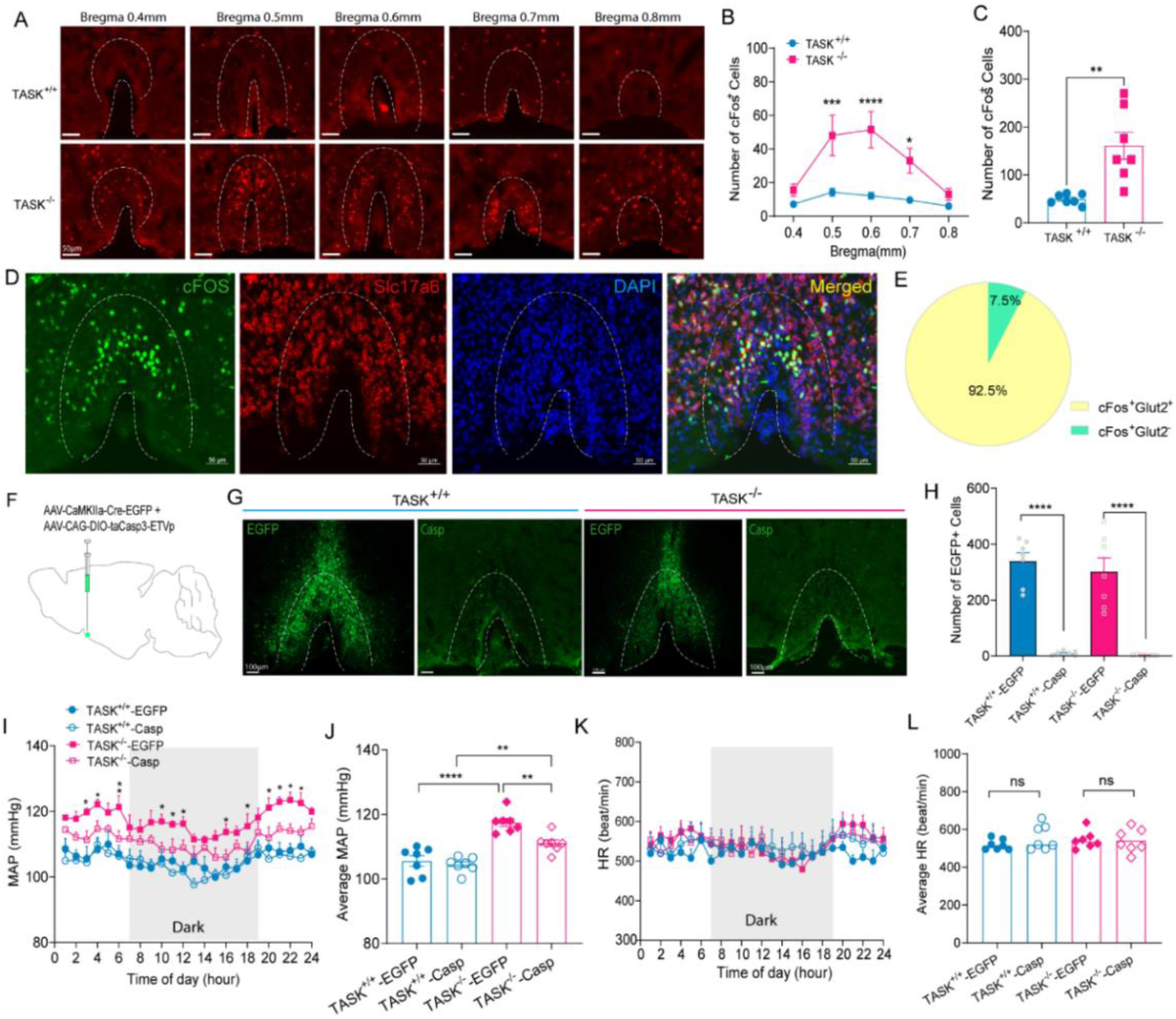
Ablation of OVLT^Glut^ neurons reduces hypertension in TASK^−/−^ Mice. (A) Images display cFos^+^ cells in OVLT at bregma levels 0.4 mm to 0.8 mm in TASK^+/+^ (top) and TASK^−/−^ (bottom) mice. Scale bar = 50 µm. OVLT regions marked by dashed lines. (B) Counts of cFos^+^ cells in OVLT at various bregma levels in two group mice (n=7 per group, two-way ANOVA with Fisher’s LSD test). (C) The total number of cFos^+^ cells come from the data presented in panel B (unpaired t test). (D) RNAscope-FISH and immunofluorescence assays to identify coexpression of cFos (green), Slc17a6 RNA (encoding vesicular glutamate transporter 2, red) and DAPI (blue) in OVLT section from TASK^−/−^ mice. Scale bars=50 μm. Pie chart showing 92.5% of cFos⁺ cells (110 cells from 3 mice) were Slc17a6⁺ (VGluT2⁺). (E) Diagram of viral injection for OVLT^Glut^ neuron ablation. (G) Validation of OVLT^Glut^ neuron ablation in mice (EGFP as control, Casp for apoptosis induction). Scale bar = 100 µm. (H) EGFP-positive neuron counts in OVLT (n=7 per group, unpaired t-test). (I) 24-hour dynamic MAP changes post-ablation in TASK^+/+^ (blue circles) and TASK^−/−^ (red squares) mice. ‘Dark’ indicates the nighttime period from 7:00 PM to 7:00 AM. n=7 per group, two-way ANOVA with Fisher’s LSD test. (J) 24-hour average MAP changes (One-way ANOVA with Tukey’s test). (K-L) 24-hour dynamic HR and average HR changes. All data are presented as mean ± SE. *P<0.05, **P <0.01, ***P <0.0005, ****P <0.0001.

To investigate whether OVLT^Glut^ neuron ablation-induced BP reduction was tied to attenuated sympathetic outflow in TASK^−/−^ mice, we assessed the effect of ganglionic blockade with hexamethonium (5 mg/kg, i.p.) during telemetric recordings. In control virus-injected mice, hexamethonium administration elicited a significantly greater BP reduction in TASK^−/−^ mice compared to TASK^+/+^ littermates (Fig.2A), confirming elevated sympathetic tone. Strikingly, this genotype-specific difference was abolished following OVLT^Glut^ neuron ablation (Fig.2B). Subgroup analysis revealed that while hexamethonium efficacy remained unchanged in OVLT^Glut^-ablated TASK^+/+^ mice, the pronounced MAP reduction observed in TASK^−/−^ controls was specifically normalized after neuroablation (Figs.2C-E). These results suggest that OVLT^Glut^ neuron ablation-induced BP reduction was tied to attenuated sympathetic outflow in TASK^−/−^ mice.

**Figure 2.**
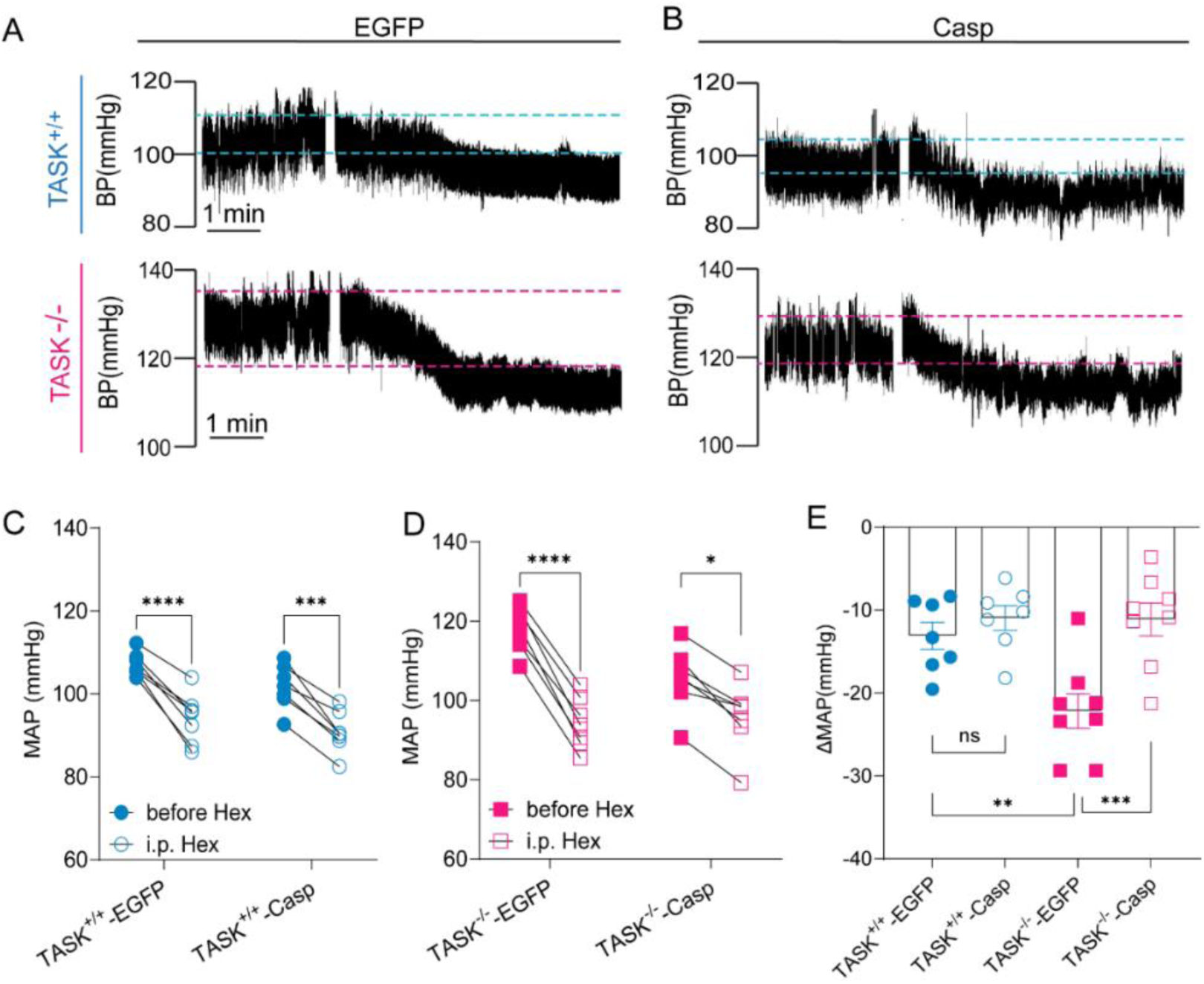
Ablation of OVLT^Glut^ neurons reduces sympathetic nerve activity in TASK^−/−^ Mice. (A) Original records of BP changes in control virus-injected TASK^+/+^ and TASK^−/−^ mice following hexamethonium (5 mg/kg, i.p.) induced ganglionic blockade. Scale bar = 1 min. The gap in the BP waveform signifies the interval for drug injection. (B) BP changes in OVLT^Glut^ neuron-ablated TASK^+/+^ and TASK^−/−^ mice following ganglionic blockade. Scale bar = 1 min. (C, D) Decrease in MAP following ganglion blockade in TASK^+/+^ (n=7 per group) and TASK^−/−^ mice (n=8 per group) with OVLT^Glut^ neuron ablation versus EGFP control (paired t-tests). (E) Peak changes in MAP in response to hexamethonium (one-way ANOVA with Tukey’s test). All data are presented as mean ± SE. *P<0.05, **P <0.01, ***P <0.0005, ****P <0.0001.

Despite significant MAP reduction, OVLT^Glut^ ablation did not fully normalize BP in TASK^−/−^ mice, suggesting additional hypertensive mechanisms. Notably, TASK^−/−^ mice exhibited lower body weight compared to age-matched TASK^+/+^ controls (Fig.S2), arguing against volume expansion as a contributing factor. It has been demonstrated that the vascular mineralocorticoid receptor (MR) pathway plays a direct role in aldosterone-mediated hypertension^46^, requiring further investigation.

### Optogenetic Activation of OVLT^Glut^ Neurons Increases BP

Recent studies have reported that activation of the OVLT can elevate sympathetic nerve activity and increase BP^28,33^. To verify this, we injected the optogenetic virus AAV-hSyn-DIO-ChR2-EYFP into the OVLT of vGlut2-Cre mice, with AAV-hSyn-DIO-EYFP as a control (Fig.3A). Four weeks post-injection, arterial BP response to laser stimulation were recorded under anesthesia. Immunofluorescence validation confirms successful expression of ChR2 in the OVLT (Fig.3B). Whole-cell patch clamp recordings showed that 1 Hz light pulses reliably evoked action potentials (APs) in OVLT^Glut^ neurons, and 5 Hz laser stimulation of their terminals in PVN slices elicited excitatory postsynaptic currents blocked by CNQX (Fig.S3). These results validate the optogenetic approach and glutamate release from the terminals. Stimulations at 1Hz, 5Hz, 10Hz, and 20 Hz rapidly elevated MAP in a pulse-rate-dependent manner (Figs.3C-F). These elevations were maintained during the stimulation and returned to baseline when the laser off (Fig.3C). Control virus-injected mice showed no BP changes in response to the same stimulations (Figs.3C-F). Additionally, HR was unaffected by stimulation across control and ChR2-expressing groups (Figs.3G-I). These findings confirm that selective OVLT^Glut^ neuron activation immediately raises arterial BP.

**Figure 3.**
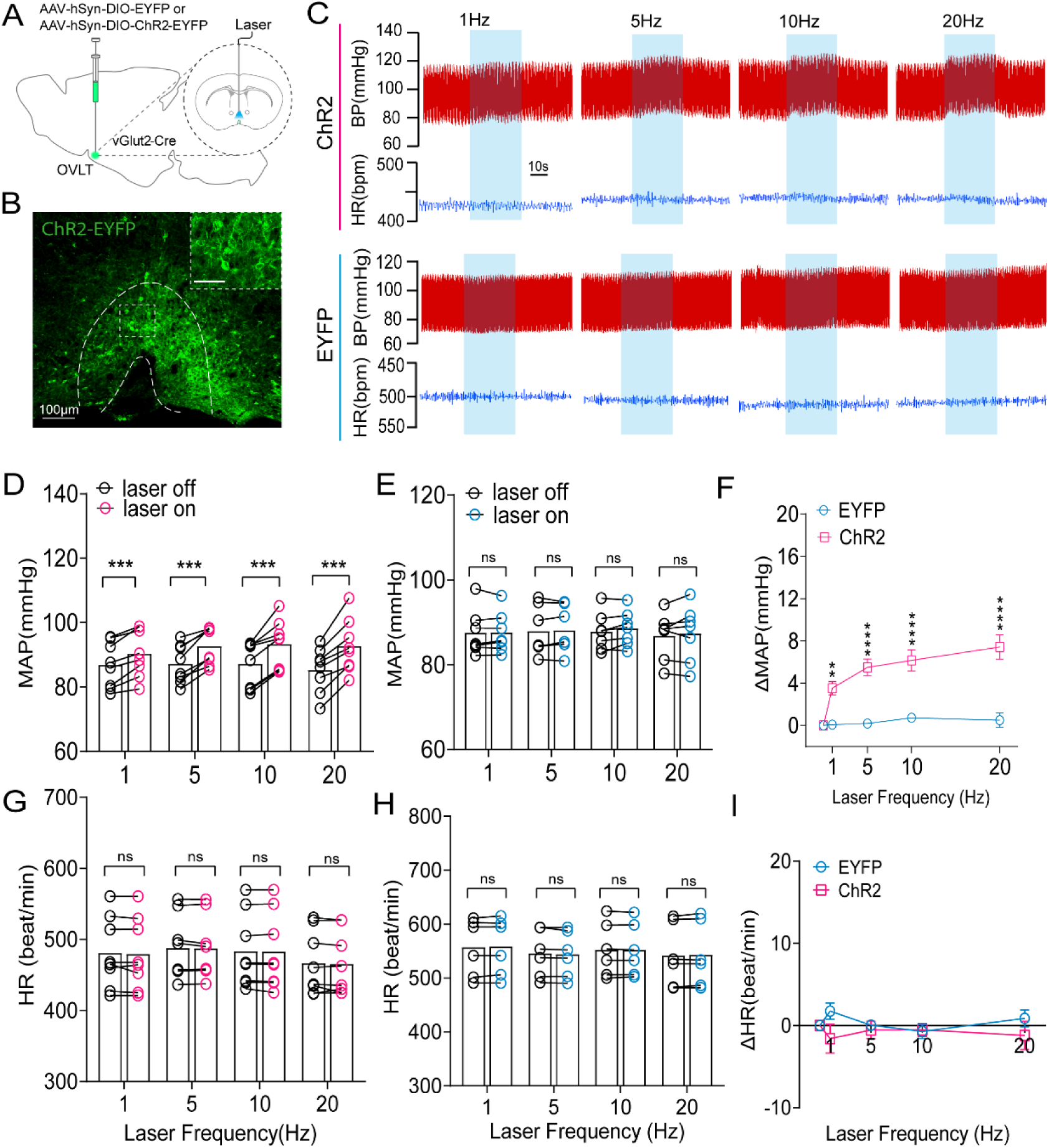
Optogenetic activation of OVLT^Glut^ neurons increases BP. (A) Schematic of optogenetics strategy in vGlut-Cre mice. (B) Immunofluorescence detection of ChR2 expression in OVLT^Glut^ neurons. Scale bar = 100 µm. Higher magnification images are in the upper right insets. Scale bar = 50 µm. (C) Representative traces of BP during OVLT^Glut^ neurons optogenetic stimulation at frequency of 1 Hz, 5 Hz, 10 Hz, and 20 Hz (473nm blue light, 20-ms pulse duration and 10 mW) in ChR2 expressing (upper panel) and control (EYFP, lower panel) group mice. Scale bar = 10 s. (D, E) BP changes with OVLT^Glut^ neurons stimulation in ChR2-expressing (red, n=8) and EYFP-expressing (blue, n=7) mice, analyzed using paired t-tests. (F) Frequency-dependent impact of optogenetic stimulation on ΔMAP (two-way ANOVA with Bonferroni post-test). (G-I) Changes in HR following optogenetic stimulation of the OVLT^Glut^ neurons at various frequencies. **P <0.01, ***P <0.0005, ****P <0.0001.

We traced the axonal projections of OVLT^Glut^ neurons and observed extensive EYFP^+^ fibers in the thalamus, hypothalamus, midbrain, and brainstem, notably in regions like the PVN and RVLM (Figs.S4-5). The PVN, a key site for presympathetic and arginine vasopressin (AVP) neurons, plays a crucial role in sympathetic nerve activity and BP regulation^42,47^. We revealed a dense network of axons from OVLT^Glut^ neurons in the PVN, including those encircling AVP neurons (Figs.4A, B), highlighting the potential regulatory role of OVLT^Glut^ neurons in vasopressin secretion. Optogenetic stimulation of these terminals induced a rapid, substantial, and reversible increase in BP, with a frequency-dependent responsiveness (Figs.4C-F). The RVLM, a pivotal sympathetic modulator, contains tyrosine hydroxylase (TH)-positive neurons that regulate spinal sympathetic pre-ganglionic neurons^48^. Our observations revealed a notable presence of OVLT^Glut^ axons in the RVLM, in close proximity to TH-positive neurons, indicating potential synaptic interactions (Figs.4G, H). Optogenetic stimulation of these terminals in the RVLM resulted in a rapid, frequency-dependent increase in BP (Figs.4I-L), similar to the response observed in the PVN. Concurrently, activation of OVLT^Glut^ axons in either the PVN or RVLM was accompanied by a slight HR decrease (data not shown), likely a baroreceptor reflex. Notably, control virus-injected mice showed no pressor response to optogenetic stimulation of OVLT^Glut^ axons in the PVN (Figs.4E, F) or RVLM (Figs.4K, L), confirming the specificity of the ChR2-induced pressor effect. In line with this, TASK^−/−^ mice exhibited elevated cFos immunoreactivity in both PVN-AVP (Fig.5A) and RVLM-TH neurons (Fig.5B) compared to controls. In addition, we performed dual-site retrograde tracing using vGlut2-Cre mice (Fig.5C). Strikingly, 40.3% of RVLM-projecting OVLT^Glut^ neurons also innervated PVN, while 21.4% of PVN-projecting neurons also innervated RVLM, revealing a potential collateral projection pattern (Fig.5D, E). Collectively, these findings establish abnormal hyperactivation of OVLT^Glut^-PVN/RVLM circuitry as a potential neuroanatomical substrate contributing to hypertension in TASK^−/−^mice.

**Figure 4.**
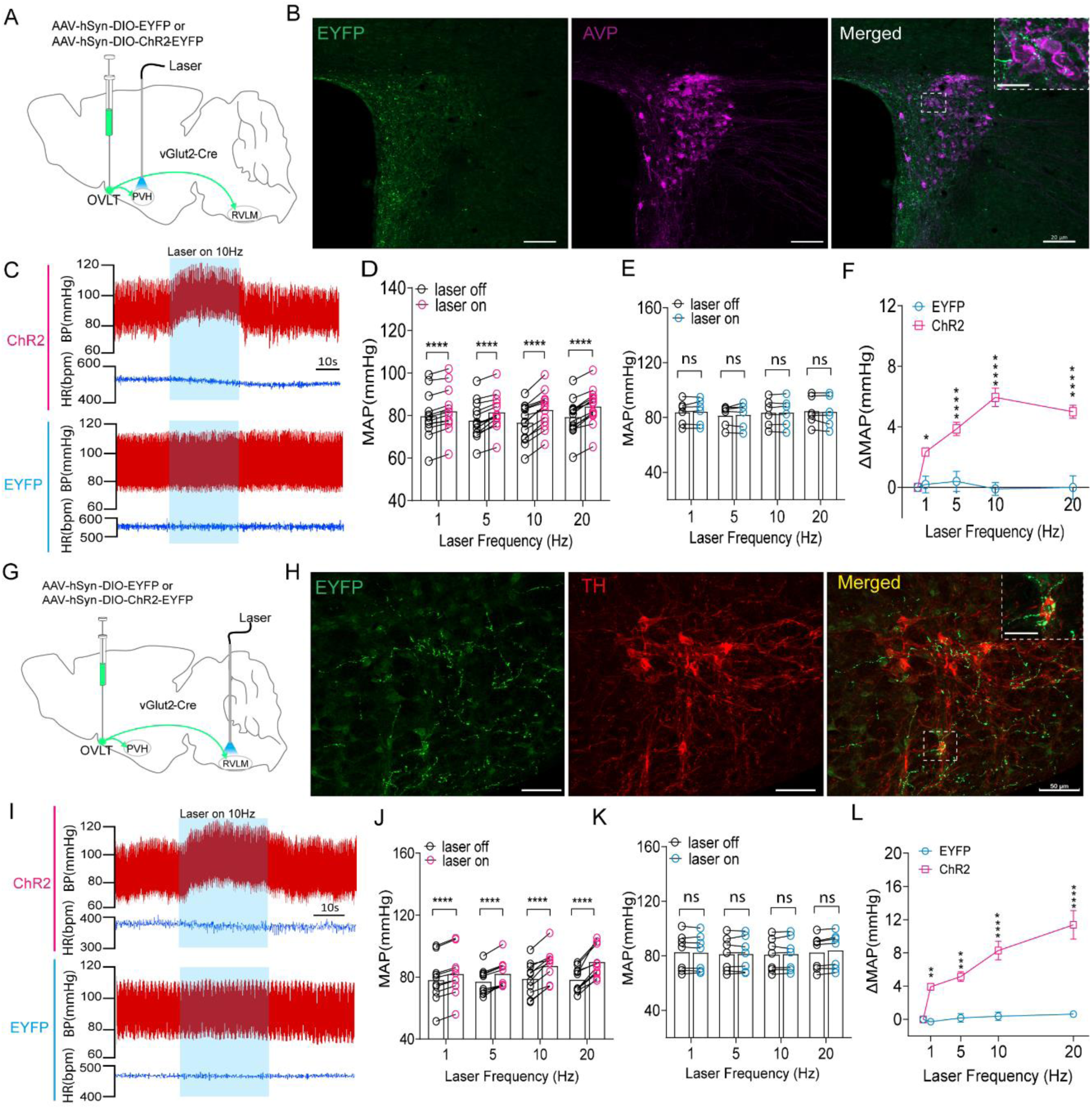
Optogenetic stimulation of OVLT^Glut^ axons in PVN and RVLM elevates BP. (A) Schematic of PVN-targeted OVLT^Glut^ axon activation in vGlut-Cre mice with ChR2-EYFP or EYFP control. (B) PVN confocal images showing EYFP^+^ axons (green), AVP immunostaining (purple), and merged view (Scale bar = 20 µm). Higher magnification images are in the upper right insets. Scale bar = 5 µm. (C) BP and HR responses to 10 Hz laser stimulation in ChR2 (top) and EYFP (bottom) mice (Scale bar = 10 s). (D, E) MAP changes post-PVN stimulation at 1 Hz, 5 Hz, 10 Hz, and 20 Hz in ChR2 (red, n=13) and EYFP (blue, n=7) mice (paired t-tests). (F) Frequency-dependent impact of optogenetic stimulation of the PVN on ΔMAP (two-way ANOVA with Bonferroni post-test). (G) Schematic of RVLM-targeted OVLT^Glut^ axon activation. (H) RVLM confocal images showing EYFP^+^ axons (green), TH immunostaining (red), and merged view (Scale bar = 50 µm). The inset shows an enlarged view of the boxed area, Scale bar = 10 µm. Images were captured as a z-stack with maximum intensity projection. (I) BP and HR variations in response to 10Hz laser stimulation in ChR2 (top) and EYFP (bottom) mice (Scale bar = 10 s). (J, K) MAP changes induced by RVLM stimulation at 1 Hz, 5 Hz, 10 Hz, and 20 Hz in ChR2 (red, n=10) and EYFP (blue, n=7) mice (paired t-tests). (L) Frequency-dependent effects of RVLM stimulation on ΔMAP (two-way ANOVA with Bonferroni post-test). *P <0.05, **P <0.01, ***P <0.0005, ****P <0.0001.

**Figure 5.**
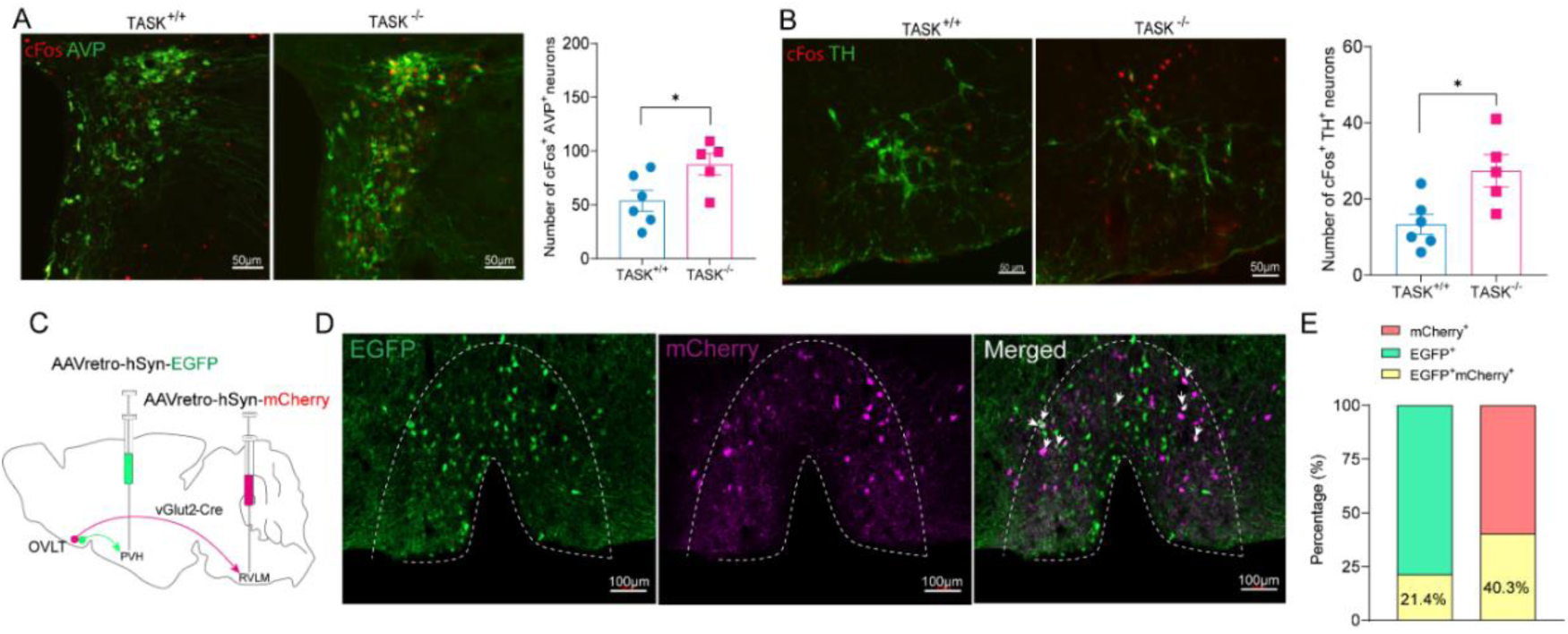
A subset of OVLT^Glut^ neurons have axonal collaterals innervating both the PVN and RVLM. (A) Representative immunofluorescence images showing increased cFos (red) in PVN-AVP neurons (green) of TASK^−/−^ mice compared to TASK^+/+^ mice. Scale bar = 50 μm. Right panel shows the quantification of co-labeled neurons (n=6 for TASK^+/+^; n=5 for TASK^−/−^, unpaired t-test, P <0.05). (B) Immunofluorescence showing increased cFos (red) in RVLM-TH neurons (green) of TASK^−/−^ vs. TASK^+/+^ mice. Scale bar = 50 μm. Right panel shows the quantification of co-labeled neurons (n=6 for TASK^+/+^; n=5 for TASK^−/−^, unpaired t-test, P <0.05.). (C) Schematic diagram of viral injection in vGlut2-Cre mice. (D) Retrogradely labeled PVN-projecting OVLT^Glut^ neurons (EGFP, green) and RVLM-projecting OVLT^Glut^ neurons (mCherry, magenta), along with their co-localization. White arrows indicate a subset of OVLT neurons that innervate both PVN and RVLM. Scale bar = 100 μm. (E) Bar graph showing the percentage of EGFP^+^ (green), mCherry^+^ (magenta), and co-labeled (yellow) neurons. Data were obtained from coronal sections of OVLT from three mice.

### Lesion of OVLT^Glut^ neurons projecting to the PVN/RVLM reduces hypertension in TASK^−/−^mice

To explore the contributions of the OVLT^Glut^-RVLM and OVLT^Glut^-PVN pathways to hypertension in TASK^−/−^ mice, we employed a Cre-dependent dual-viral strategy to target these circuits (Figs. 6A, H). AAVretro-CaMKIIa-Cre-GFP was injected into either RVLM or PVN to retrogradely label and genetically target OVLT^Glut^ neurons projecting to these regions. Subsequent OVLT injection of AAV-CAG-DIO-taCasp3-ETVp induced caspase-3-mediated apoptosis specifically in Cre-expressing neurons, with AAV-CAG-DIO-GFP serving as control. In control mice, GFP-labeled neurons were mainly resided inside the OVLT, with OVLT^Glut^-PVN neurons located medially near the ventricles and OVLT^Glut^-RVLM neurons showing a scattered distribution (Figs.6B, I). In contrast, mice with OVLT^Glut^ neuron ablation showed few GFP-positive neurons, indicating successful targeting (Figs.6C, J).

**Figure 6.**
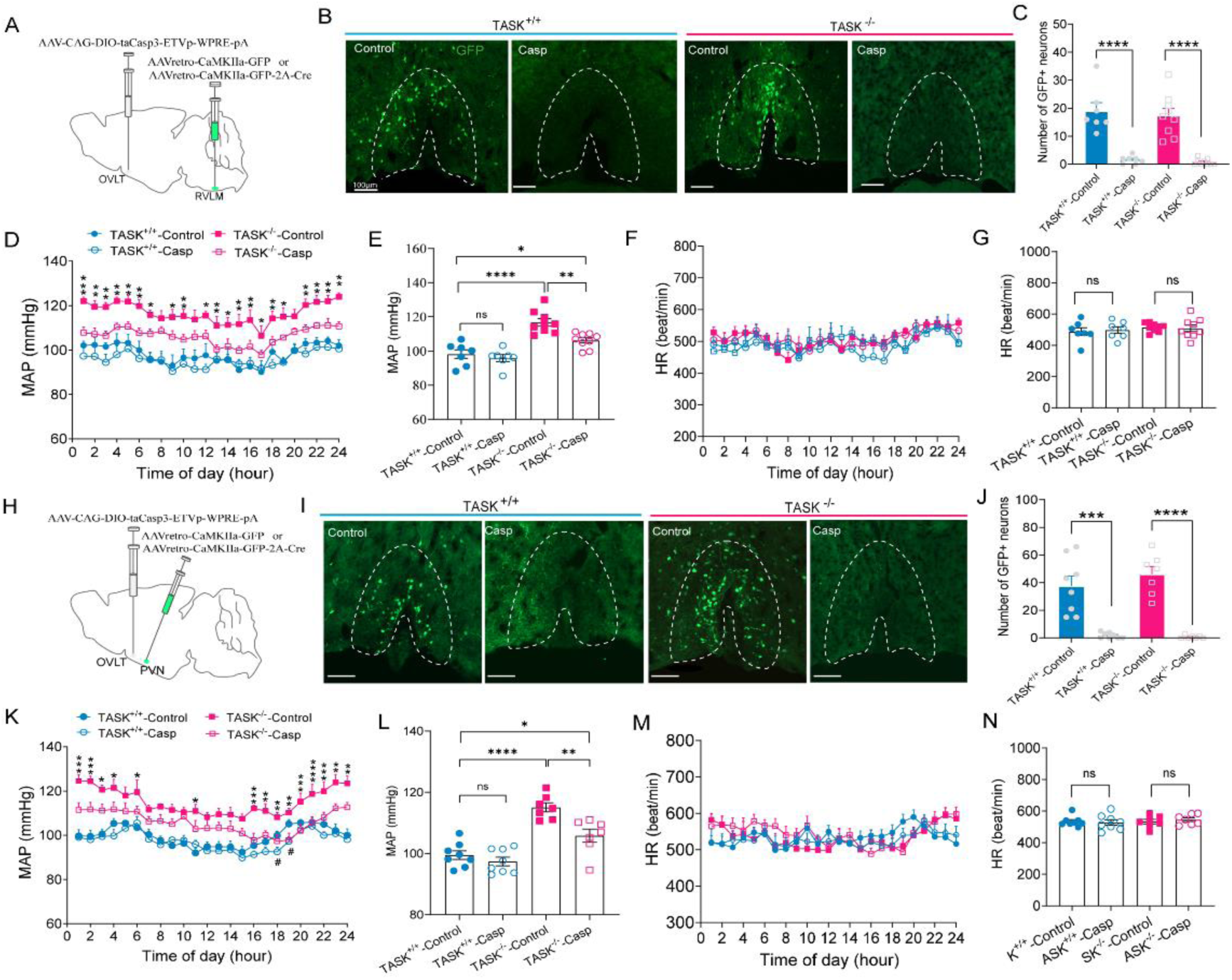
Ablation of OVLT^Glut^-PVN/RVLM neurons reduces hypertension in TASK^−/−^ mice. (A) Schematic of virus injection for OVLT^Glut^-RVLM neuron ablation. (B) OVLT^Glut^-RVLM neuron images in TASK^+/+^ and TASK^−/−^ mice (Casp for apoptosis, Control for non-apoptotic). Scale bar = 100 μm. (C) Four weeks post-injection, GFP^+^ neuron count significantly decreased in the apoptosis group vs. control in both strains (TASK^+/+^: n=7 for both groups; TASK^−/−^: n=9 for both groups). (D) Ablation of OVLT^Glut^-RVLM neurons significantly reduced MAP at most time points over a 24-hour period in TASK^−/−^ mice (red), with no significant effect in TASK^+/+^ mice (blue). (E) Average 24-hour MAP summary. (F, G) Effects on 24-hour HR dynamics and average HR in both strains. (H) Schematic of virus injection for OVLT^Glut^-PVN neuron ablation. (I, J) PVN-projecting OVLT^Glut^ neuron images in each group. Scale bar = 100 μm. (K, L) OVLT^Glut^-PVN neuron ablation impacts on 24-hour MAP dynamics and average MAP in both mouse strains (Casp vs. Control, TASK^+/+^: n=8 for both groups; TASK^−/−^: n=7 for both groups). (M, N) Effects on 24-hour HR dynamics and average HR. Dynamic BP and HR data over 24 hours were analyzed using two-way ANOVA with Fisher’s LSD post-test, and other group comparisons used unpaired t-test. Scale bars = 100 µm. TASK^−/−^-Control vs. TASK^−/−^-Casp, *P <0.05, **P <0.01, ***P <0.0005, ****P <0.0001; TASK^+/+^-Control vs. TASK^+/+^-Casp, ^#^P <0.05; ns, no significance.

In TASK^−/−^ mice, selective ablation of OVLT^Glut^-RVLM neurons led to significant reductions in MAP at certain times within 24 hours, while TASK^+/+^ littermates showed no such changes, highlighting genotype-specific effects (Fig.6D). Accordingly, the 24-hour average MAP in TASK^−/−^ mice was significantly reduced yet remained higher than the baseline levels observed in TASK^+/+^ mice (11.67±2.2 *vs.* 106.2±1.5 *vs.* 98.2±2.5, Fig.6E). The HR remained unchanged in both genotypes post-ablation (Figs.6F, G). We employed a similar apoptotic viral strategy to ablate the OVLT^Glut^-PVN pathway (Fig.6H-J) and found results similar to those obtained with the OVLT^Glut^-RVLM pathway ablation (Figs.6K-N). The results imply that aberrant activation of either the OVLT^Glut^-PVN pathway, the OVLT^Glut^-RVLM pathway, or both, might underlie the hypertension observed in TASK^−/−^ mice.

### Elevated Excitability of OVLT^Glut^-PVN/RVLM Neurons in TASK^−/−^ Mice

Building on the hypertension alleviation in TASK^−/−^ mice post-ablation of OVLT^Glut^-PVN/RVLM neurons, we next examined potential intrinsic functional changes. We conducted whole-cell current clamp recordings from acute OVLT slices of TASK^+/+^ and TASK^−/−^ mice to assess excitability and firing properties of these neurons. OVLT^Glut^-PVN/RVLM neurons were retrogradely labeled via AAVretro-CaMKIIa-GFP three weeks prior. The RMP was measured at I=0 in current clamp immediately after membrane rupture. Neuronal excitability was evaluated by applying 500-ms current step injections.

We found that OVLT^Glut^-PVN/RVLM neurons exhibited a significantly depolarized RMP in TASK^−/−^ mice compared to TASK^+/+^ mice (−56.8±0.9 mV *vs.* −62.3±0.9 mV) and increased AP firing upon current injection, with no changes in input resistance and membrane capacitance, suggesting enhanced excitability (Figs.7A-E). Mechanistically, while hyperaldosteronism and sodium imbalance may contribute to neuronal hyperexcitability, the absence of TASK-1/TASK-3 channels also likely plays a primary role, as these Two-Pore Domain Potassium Channels are critical for RMP stabilization and excitability regulation. Single-cell sequencing confirms robust expression of TASK-1 and TASK-3 in both glutamatergic and GABAergic neurons of the OVLT (Fig.7N). Notably, MRs are also highly expressed in the OVLT (Fig.S7). To evaluate the role of excessive aldosterone, we treated TASK^−/−^ mice with the MRs antagonist spironolactone (100 mg/kg, once per day for 7 days by oral gavage). This intervention restored the AP firing frequency to levels comparable to TASK^+/+^ mice (Fig.7C), although the RMP remained slightly depolarized (Fig.7A). This evidence supports the notion that excessive aldosterone signaling promotes neuronal hyperexcitability, while the possibility of compensatory adaptations due to TASK channel loss remains a consideration.

**Figure 7.**
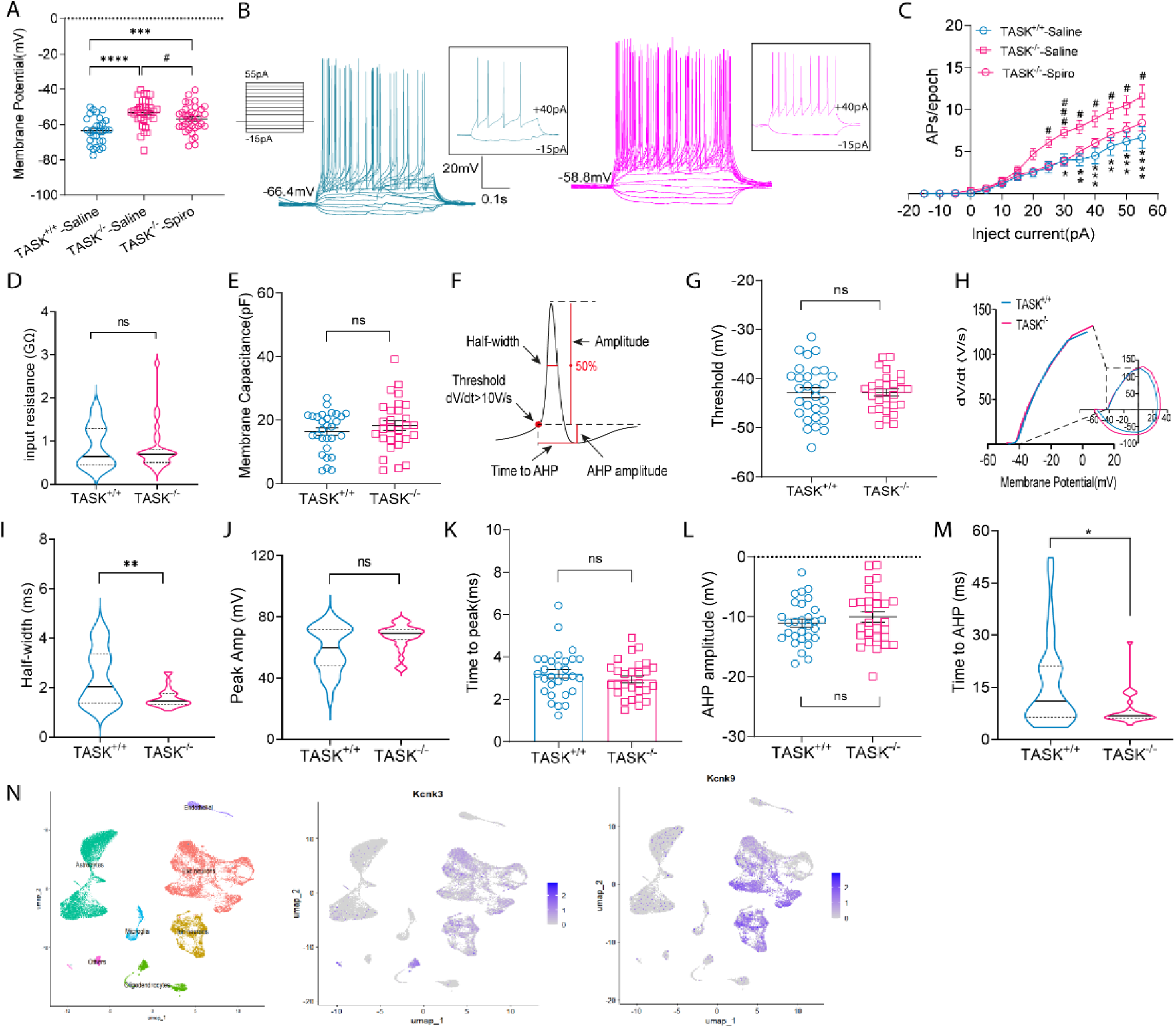
OVLT^Glut^-PVN/RVLM neurons exhibit increased excitability in TASK^−/−^ mice. (A) The RMP of OVLT^Glut^-PVN/RVLM neurons in TASK^+/+^ mice (n=29 neurons) and TASK^−/−^ mice (n=34 neurons) gavage saline, and TASK^−/−^ mice (n=36 neurons) gavage spironolactone (Spiro) for 7 days. Data analyzed by one-way ANOVA with Fisher’s LSD test. *P, TASK^−/−^-Saline vs. TASK^+/+^-Saline; ^#^P, TASK^−/−^-Saline vs. TASK^−/−^-Spiro. (B) Representative current-clamp recordings of OVLT^Glut^-PVN/ RVLM neurons from TASK^+/+^ (blue) and TASK^−/−^ (red) mice. Inset: representative trace in response to +40 pA and −15 pA injection. (C) Action potential (AP) numbers of OVLT^Glut^-PVN/RVLM neurons evoked by step current injection in TASK^+/+^-Saline (n=29 neurons), TASK^−/−^-Saline (n=28 neurons), and TASK^−/−^-Spiro (n=23 neurons) mice. Data analyzed by two-way ANOVA with Bonferroni test. *P, TASK^−/−^-Saline vs. TASK^+/+^-Saline; ^#^P, TASK^−/−^-Saline vs. TASK^−/−^-Spiro. (D, E) Input resistance and membrane capacitance values. (F) AP characteristics measurement. AHP, after hyperpolarization. (G, H) Voltage threshold and associated phase-plane plots. (I-M) AP half-width, peak amplitude, time to peak, AHP amplitude, and time to AHP values of OVLT^Glut^-PVN/ RVLM neurons in TASK^+/+^ and TASK^−/−^ mice. (N) UMAP of OVLT cell clustering in wild type mice and UMAP of Kcnk3 and Kcnk9 gene distributions. Pairwise comparisons were estimated using Mann–Whitney tests or unpaired t test. ^#^P <0.05, *P <0.05, **P <0.01, ***P <0.0005, ****P <0.0001.

We further compared the AP properties between TASK^+/+^ and TASK^−/−^ mice by quantifying the threshold, amplitude, half-width, AHP amplitude, and time to AHP from the single AP evoked by the rheobase current^41^ (Fig.7F). Intriguingly, we observed a decrease in both the AP half-width (Fig.7I) and time to AHP (Fig.7M), while the AP threshold (Figs.7G, H), amplitude (Fig.7J), time to peak (Fig.7K), and AHP amplitude (Fig.7L) remained unchanged. These findings further reinforce the notion of enhanced OVLT^Glut^-PVN/RVLM neurons excitability in TASK^−/−^ mice, as the reduced AHP time and AP half-width suggest a more shortened repolarization phase, which could contribute to the higher number of APs fired in response to current injection.

### Chronic Aldosterone Treatment Enhances OVLT^Glut^-PVN/RVLM Neurons Excitability

To elucidate the specific impact of excessive aldosterone, TASK^+/+^ mice received subcutaneous osmotic pumps delivering aldosterone (900 μg/mL), while controls received vehicle (5% ethanol). OVLT^Glut^-PVN/RVLM neurons were visualized by labeling them retrogradely using AAVretro-CaMKIIa-GFP, and electrophysiological studies were conducted 7 days post-implantation (Fig.8A). We observed a significant depolarization of approximately 3 mV in the RMP of OVLT^Glut^-PVN/RVLM neurons in aldosterone-treated mice compared to the vehicle group (62.4±0.9 mV *vs.* 59.2±0.8 mV, Fig.8B). Under current-clamp mode, the number of AP evoked by injected current was notably increased (Figs.8C, D), while the input resistance (Fig.8E) and membrane capacitance (Fig.8F) without significant differences. This indicates that chronic aldosterone exposure enhances the excitability of OVLT^Glut^-PVN/RVLM neurons. Subsequently, we analyzed the characteristics of APs elicited by the rheobase current and found that the AP half-time (Fig.8H), and time to AHP (Fig.8L) were shortened, while the threshold (Fig.8G), amplitude, peak time, and AHP amplitude showed no significant changes (Figs.8I-K). These findings closely resemble those in TASK^−/−^ mice, suggesting that the depolarization of the RMP and the increased neuronal excitability in OVLT^Glut^-PVN/RVLM neurons are likely driven, at least in part, by excessive aldosterone. Nevertheless, the role of aldosterone in this process still lacks compelling evidence and remains to be investigated in mice with a zona glomerulosa-selective deletion of TASK-1 and TASK-3 channels^49^.

**Figure 8.**
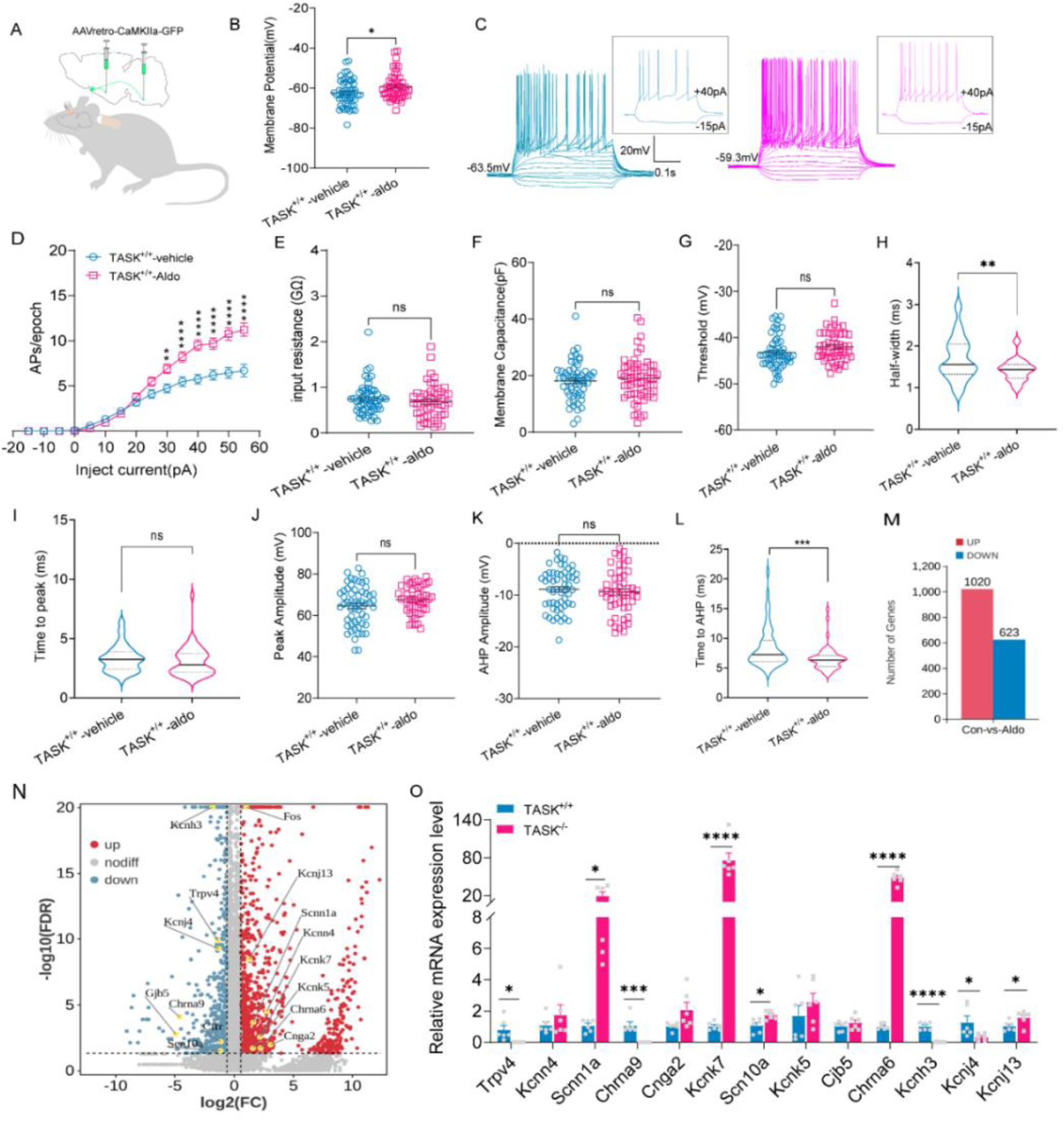
Aldosterone enhances excitability of OVLT^Glut^-PVN/RVLM neurons in TASK^+/+^ mice. (A) Schematic of viral strategy for labeling OVLT^Glut^-PVN/RVLM neurons in TASK^+/+^ mice, with osmotic minipump and intracerebroventricular injection catheter. (B) The RMP of OVLT^Glut^-PVN/RVLM neurons comparison between vehicle-treated mice (n=56 neurons) and aldosterone-treated (n=50 neurons, unpaired t-test). (C) Representative current-clamp recordings of OVLT^Glut^-PVN/ RVLM neurons from vehicle (blue) and aldosterone (red) treated mice. Inset: response to +40 pA and −15 pA current injections. (D) AP numbers of OVLT^Glut^-PVN/RVLM neurons evoked by step current injection from vehicle-treated (n = 56 neurons) and aldosterone-treated mice (n = 50 neurons, two-way ANOVA with Bonferroni post-test). (E-L) Comparative analysis of input resistance, membrane capacitance, threshold, half-width, time to peak, peak amplitude, AHP amplitude, and time to AHP between vehicle-treated (n=56 neurons) and aldosterone-treated (n=50 neurons) mice. (M) RNA sequencing showing gene expression changes in aldosterone-treated vs. vehicle-treated mice. (N) Volcano plot of genes significantly upregulated (red) and downregulated (blue) in aldosterone-treated (n=6) mice compared to vehicle (n=4) group. The cut-off of false discovery rate (FDR)-adjusted P < 0.05 and the absolute logarithmically transformed fold-change (|FC|) ≥ 1.5. (O) qPCR validation of ion channel gene expression in OVLT of TASK^+/+^ and TASK^−/−^ mice (n=6 in each group). Data are presented as the mean ± SEM from three independent biological replicates. Pairwise comparisons were estimated using Mann– Whitney tests or unpaired t-test. *P <0.05, **P <0.01, ***P <0.0005, ****P <0.0001.

### The Potential Molecular Mechanisms Underlying the Enhancement of Neuronal Excitability by Chronic Aldosterone administration

To explore the molecular basis of heightened neuronal excitability, RNA-seq was performed on mice with aldosterone- or vehicle-infused pumps over 7 days. RNA-seq analysis revealed significant transcriptional alterations in the OVLT of aldosterone-treated TASK^−/−^ mice compared to vehicle-treated controls (Fig.8M). Specifically, 1,020 genes were upregulated and 623 genes were downregulated in the OVLT of aldosterone-exposed mice, indicating widespread transcriptomic reprogramming in this hypothalamic nucleus. Consistent with these findings, a marked increase in cFos gene expression, a marker of neuronal activation, in the OVLT of aldosterone-treated mice (Fig.8N).

Considering the crucial role of ion channels in the regulation of neuronal excitability, we focused on the mRNA expression of ion channels as defined by IUPHAR^50^ (Table S3). Among these differentially expressed genes, we identified 16 ion channel genes (Fig.8N, Table S4). The sodium channels included the epithelial sodium channel (ENaC, Scnn1a) and the voltage-gated sodium channel Nav1.8 (Scn10a). For potassium channels, we observed variations in inwardly rectifying potassium channels Kir2.3 (Kcnj4) and Kir7.1 (Kcnj13), two-pore domain potassium channels TASK2 (Kcnk5) and TWIK3 (Kcnk7), the intermediate conductance calcium-activated potassium channel KCa3.1 (Kcnn4), and the voltage-gated potassium channel Kv12.2 (Kcnh3). Besides, there are the transient receptor potential cation channel TRPV4 (Trpv4) and the cyclic nucleotide-gated channel CNGA2 (Cnga2), as well as other genes such as Gjb5, Gjb6, Cftr, Chrna1, Chrna6, and Chrna9, which encode gap junction proteins and other receptor proteins (Fig.8N). These aldosterone-induced ion channel genes might serve as the molecular basis for the changes in the electrophysiological characteristics of OVLT^Glut^-PVN/RVLM neurons. Importantly, qPCR results confirmed that the expression of Trpv4, Scnn1a, Scn10a, Chrna9, Chrna6, Kcnh3, Kcnj4, Kcnk7, and Kcnj13 exhibited similar alterations in TASK^−/−^ mice compared to TASK^+/+^ mice (Fig.8O). The altered expression of Scnn1a, Scn10a, Chrna6, and Kcnh3 in the OVLT of TASK^−/−^ mice can be reversed by spironolactone treatment (Fig.S6). This coincidence leads us to hypothesize that the alterations in these ion channel genes in TASK^−/−^ mice are likely associated with their excessive aldosterone, although this inference still lacks direct evidence. The incomplete knowledge of the genes selectively targeted by aldosterone/MR, especially in the brain, adds a challenge to elucidating the cause of the aberrant expression of these genes in TASK^−/−^ mice^51^. 11β-HSD_2_ is key for aldosterone’s genomic action via MR, while GPER1 is linked to its nongenomic effects^52^. However, both Hsd11b2 and Gper1 are undetectable in the OVLT (Fig.S7), suggesting that different mechanisms or pathways may be at play.

### Aldosterone amplifies Central Pressor Effect of Salt

To investigate whether chronic aldosterone treatment could elevate arterial BP in mice, given its effect on enhancing the excitability of OVLT^Glut^-PVN/RVLM neurons, we conducted a series of experiments. Notably, 7 days aldosterone administered alone or drinking 1% NaCl solution alone did not significantly alter MAP in TASK^+/+^ mice (Fig.9A). However, a significant increase in MAP was observed in mice subjected to a combination of aldosterone treatment and 1% NaCl solution intake (Fig.9A). Consistently, the cFos expression in OVLT^Glut^-PVN/RVLM neurons, labeled using the same method as described above, was significantly higher in mice from the aldosterone/1% NaCl combination treatment group compared to the other groups (Figs.9B, C). This suggests that aldosterone may elevate BP by sensitizing the central pressor effect of sodium. To further validate this hypothesis, we performed intracerebroventricular infusions of a hypertonic solution (450 mM [Na^+^]) at a rate of 0.4 μL/min for 10 minutes via a pre-implanted cannula in anesthetized mice (Fig.8A), while continuously monitoring BP. The hypertonic infusion elicited a progressive rise in BP in both aldosterone- and vehicle-treated group mice injected with the control virus (Figs.9D-E). Crucially, the magnitude of this BP increase was significantly greater in the aldosterone-treated mice compared to the vehicle controls (Fig.9F). This finding underscores that aldosterone enhances the central pressor response to salt, indicating a synergistic interaction that contributes to BP elevation. Furthermore, when OVLT^Glut^-PVN/RVLM neurons were ablated (Fig.9D), the observed difference in BP response induced by hypertonic solution infusion between the aldosterone-treated and control groups was abolished (Figs.9G-H). Taken together, our results indicate that aldosterone treatment induces central neuroplastic changes in the OVLT^Glut^-PVN/RVLM neurons, thereby sensitizing the hypertensive response to salt.

**Figure 9.**
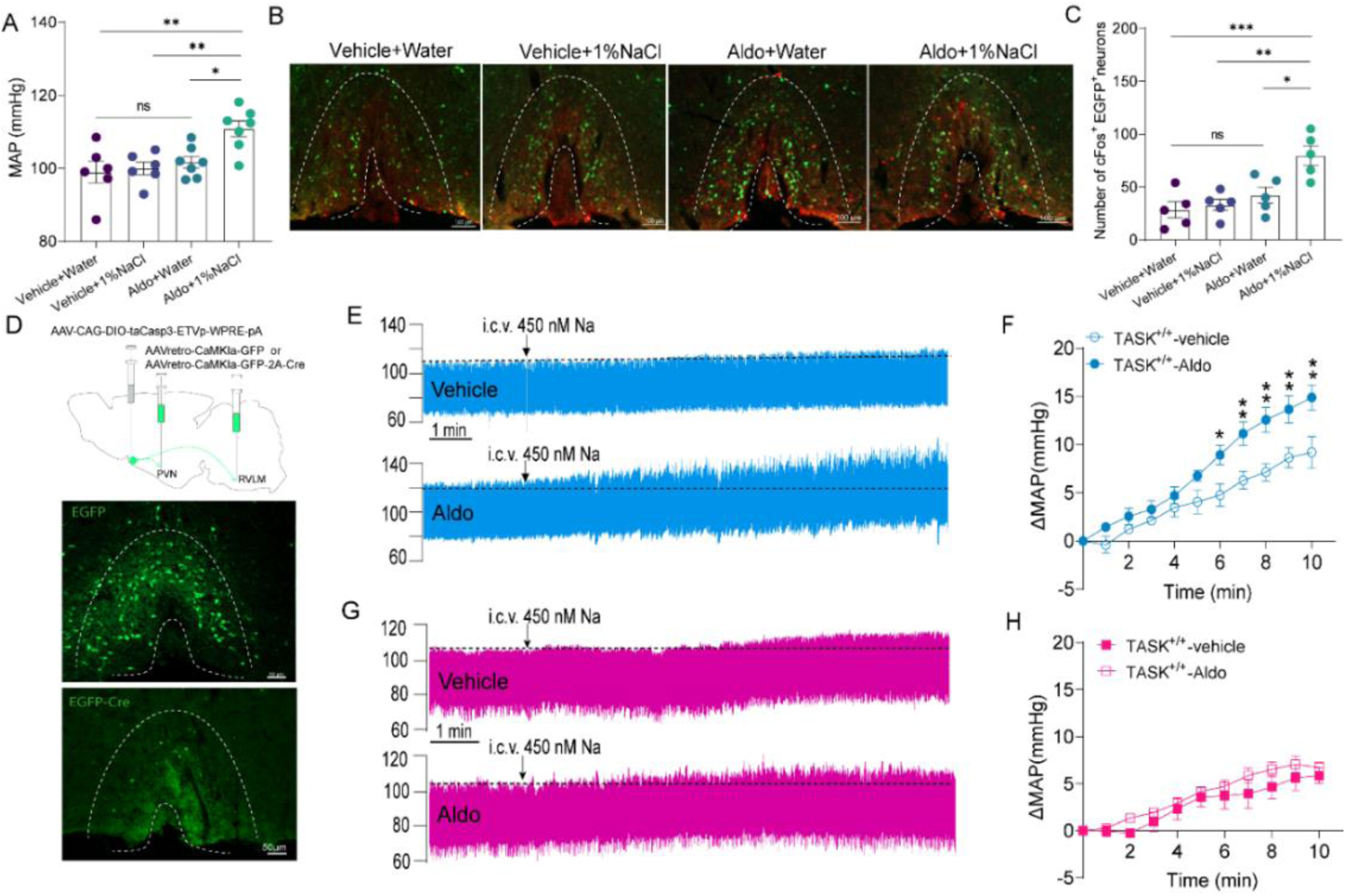
OVLT^Glut^-PVN/RVLM neurons contribute to aldosterone-sensitized central sodium pressor responses. (A) Mean arterial pressure (MAP) in TASK^+/+^ mice chronically treated with aldosterone or vehicle and given either water or 1% NaCl solution for 7 days. Sample sizes: n=6 in both vehicle-treated groups, and n=7 in both aldosterone-treated groups. Data analyzed by one-way ANOVA with Tukey’s multiple comparisons test. (B) Representative immunofluorescence images showing cFos expression in OVLT^Glut^-PVN/RVLM neurons across groups. Scale bar = 100 μm. (C) Quantification of cFos expression. n=5 per group. One-way ANOVA with Tukey’s test. (D)Viral strategy for OVLT^Glut^-PVN/RVLM neuron ablation and ablation efficiency images. Scale bar = 50 μm. (E) Original records of BP changes in response to intracerebroventricular (i.c.v.) injection of hypertonic solution (450 nM Na^+^ aCSF solution) in control virus injected mice treated with aldosterone or vehicle. Scale bar = 1 min. (F) Statistical analysis of MAP changes over 10 minutes. (G) Original records of BP changes in response to hypertonic solution in OVLT^Glut^-PVN/RVLM neuron-ablated mice treated with aldosterone or vehicle. Scale bar = 1 min. (H) Statistical analysis of MAP changes over 10 minutes in OVLT^Glut^-PVN/RVLM neuron-ablated mice treated with aldosterone or vehicle. For panels F and H, the data were analyzed using two-way ANOVA with Bonferroni’s multiple comparisons test (n=5 in each group). *P <0.05, **P <0.01, ***P <0.0005.

## Discussion

In this study, we identify a pivotal role of OVLT^Glut^ neurons in hyperaldosteronism-associated hypertension in TASK^−/−^ mice. Our findings highlight the enhanced excitability of OVLT^Glut^ neurons contributes to hypertension in TASK^−/−^ mice through regulation of OVLT^Glut^-PVN/RVLM circuits. The hyperexcitability of these neurons is partly due to aldosterone-induced changes in ion channel expression spectrum, contribute to hypertension by amplifying central sodium sensitivity.

### The overexcitation of OVLT^Glut^ neurons is implicated in the hypertension of TASK^−/−^ mice

The OVLT, with ∼80% glutamatergic neurons and an incomplete blood-brain barrier, senses and integrates blood-borne signals to coordinate physiological responses^44^. Recent in vivo single-unit recordings by Sean D. Stocker indicate that elevated neuronal discharge of NaCl-sensitive OVLT neurons contributes to hypertension in Dahl S rats, characterized by heightened baseline activity and exaggerated responses to hypertonic NaCl^33^. This evidence indicates that OVLT neuronal plasticity changes are capable of causing salt-sensitive hypertension. Electrical or optogenetic stimulation of OVLT neurons increases sympathetic nerve activity and arterial BP^28^. Importantly, chronic chemogenetic activation of OVLT neurons produces hypertension associated with enhanced sympathetic outflow^28^. Collectively, these observations raise the hypothesis that hyperexcitability of OVLT salt-sensitive neurons leads to enhanced sympathetic nerve activity and elevated BP.

In our study using TASK^−/−^ mice (a well-defined PA model), we found increased cFos expression in OVLT neurons, with 92.5% of these being glutamatergic. Optogenetic activation of OVLT^Glut^ neurons in vGlut2-Cre mice rapidly increased BP in a frequency-dependent manner. Selective ablation of OVLT^Glut^ neurons significantly reduced hypertension and eliminate the greater depressor responses to ganglionic blockade in TASK^−/−^ mice, with minimal effects on TASK^+/+^ mice. This suggests that overexcitation of OVLT^Glut^ neurons contributes to sustained hypertension in TASK^−/−^ mice, likely through enhanced sympathetic activity. Our findings are corroborated by other studies, which have demonstrated that selective ablation of the OVLT prevents or reverses hypertension in several hypertension models, including the salt-sensitive Dahl model^53^ and the DOCA-salt model^54^. It is worth noting that the ablation of OVLT^Glut^ neurons did not fully reverse the hypertensive state, indicating the involvement of additional mechanisms.

One possibility is the SFO, another central osmoreceptor and key sodium-sensing site, which also regulates sympathetic activity and BP. Although we did not specifically investigate the role of the SFO in this study, it is plausible that it may also be implicated in the hypertension observed in TASK^−/−^ mice. Nomura K et al. have demonstrated that selective lesions of the OVLT, in contrast to lesions of the SFO, abolish the BP response to salt^42^. This finding suggests that the OVLT may be more sensitive to sodium, while the SFO may play a more prominent role in AngII-mediated BP regulation. The MnPO, which is in close proximity to the OVLT and contains glutamatergic neurons, receives inputs from both the OVLT and the SFO ^55,56^ and may also be implicated in the hypertensive phenotype observed in TASK^−/−^ mice. Despite the potential for aldosterone to contribute to hypertension through directly vasoconstrictive and promote renal sodium retention mechanisms, we observed that TASK^−/−^ mice exhibited lower body weight compared to their TASK^+/+^ counterparts. This finding suggests that the expansion of blood volume may not be a major contributor to hypertension in TASK^−/−^ mice. However, it is possible that elevated aldosterone levels may increase metabolic rate^57^, which could account for the reduced body weight in these mice. Additionally, TASK-1 or TASK-3 channels inhibition in peripheral arterial chemoreceptors is known to activate sympathetic nerve activity in response to decreased oxygen partial pressure or acidosis^58^.

Clinical and experimental studies have shown that excessive aldosterone heightened salt sensitivity, a key factor in salt-sensitive hypertension and PA-related hypertension^13,34,36,43^. Consistent with this evidence, our research has further established that both a low-sodium diet and intracerebroventricularly administration of spironolactone can effectively mitigate hypertension in TASK^−/−^ mice^49, 50^. Landmark studies by Alan Kim Johnson et al. indicate that subpressor doses of aldosterone or AngII administered intracerebroventricularly can induce salt sensitivity in normotensive models, effectively priming central circuits to hyperrespond to NaCl challenges^4,8^. Makhanova et al. further suggest that a moderate increase in aldosterone synthase expression can enhance BP responsiveness to salt^11^. Additionally, we found that aldosterone exposure alone does not significantly increase BP but potentiates the central pressor effects of intracerebroventricular hypertonic solution infusion, an effect that is abolished by OVLT^Glut^ neuron lesioning. Given these findings, we speculate that the excessive aldosterone synergistic with high sodium context to enhance sympathetic activity is a crucial contributor to the hypertension observed in TASK^−/−^ mice.

### OVLT^Glut^-PVN/RVLM Circuit Mediates Hypertension in TASK^−/−^ Mice

Though prior studies have shown that OVLT neurons activated by high sodium relay excitement directly or indirectly to sympathetic control centers like the PVN^59^ and RVLM^32,60^, the complete brain projections of OVLT^Glut^ neurons are still not fully understood. Our study indicates they project widely to the thalamus, hypothalamus, brainstem, including the PVN and RVLM. While direct projections from OVLT^Glut^ neurons to the PVN are certain^21^ and PVN to RVLM projections are confirmed^61^, to our knowledge, no direct axonal projections from OVLT to RVLM have been documented. Sean D. Stocker showed that OVLT neurons enable increased dietary salt to enhance RVLM sympathetic neuron responsiveness, yet without direct anatomical link proof ^32,60^. We have evidence supporting direct OVLT^Glut^ to RVLM projections: (1) Anterograde tracing found OVLT^Glut^ axonal terminals in RVLM, likely not pass-through fibers as they tightly encircle TH^+^ neurons (primary presympathetic neuron subtype). (2) Retrograde labeling from RVLM identified labeled OVLT^Glut^ neurons, some near the ventricular area, reinforcing the direct projection. (3) Optogenetic activation of the RVLM terminals originating from OVLT^Glut^ neurons resulted in an acute increase BP. (4) Genetic ablation of the projections from OVLT^Glut^ neurons to the RVLM led to a significantly reduction in BP in TASK^−/−^ mice, but not in TASK^+/+^ mice.

The PVN is a key hypothalamic nucleus regulating BP and fluid balance. It contains presympathetic neurons modulating sympathetic outflow and osmosensitive vasopressinergic neurons releasing vasopressin to maintain water homeostasis via pituitary secretion. Indeed, studies have reported that AVP plays a primary role in DOCA-salt hypertension by enhancing vascular reactivity and via central mechanisms requiring OVLT region integrity^29,30^. Our experimental results indicate that OVLT^Glut^ neurons innervate PVN^AVP^ neurons. In TASK^−/−^ mice, cFos expression in PVN^AVP^ and RVLM^TH^ neurons was markedly increased compared to TASK^+/+^ mice. Selectively activating OVLT^Glut^ neuron terminals projecting to the PVN rapidly elevated BP, while lesioning the OVLT^Glut^-PVN pathway significantly reduced hypertension in TASK^−/−^ mice. This suggests that both the OVLT^Glut^-PVN and OVLT^Glut^-RVLM pathways contribute to hypertension in TASK^−/−^ mice. However, the distinct roles of these pathways were not further explored. It is plausible that a subset of OVLT^Glut^ neurons may have axonal collaterals innervating both the PVN and RVLM. Indeed, we found that 40.3% of RVLM-projecting OVLT^Glut^ neurons also innervate the PVN, and 21.4% of PVN-projecting OVLT^Glut^ neurons also target the RVLM. Additionally, we cannot rule out that stimulating OVLT^Glut^ axons terminal in the PVN might activate OVLT neurons projecting to the RVLM to increase BP. We also acknowledge that other pathways beyond the OVLT^Glut^-PVN/RVLM may contribute to hypertension in TASK^−/−^ mice. OVLT^Glut^ neurons project to multiple nuclei in the thalamus, hypothalamus, and midbrain, some of which are known to regulate BP^62^.

### The possible molecular mechanisms behind the increased excitability of OVLT^Glut^-PVN/RVLM neurons

Our present patch-clamp data revealed that OVLT^Glut^-PVN/RVLM neurons exhibited increased excitability in TASK^−/−^ mice, which was characterized by depolarization of the RMP, higher frequency of AP firing in response to current injection, as well as shortened AP half-time and repolarization times. Depolarization of the RMP while the threshold potential remains unchanged implies that the neuron becomes more easily excitable and more likely to trigger an AP. Narrowing the AP half-width and shortening the repolarization phase can significantly enhance neuronal excitability by allowing a quicker return to threshold potential, reducing the refractory period, and enabling rapid successive firings^63^. The heightened excitability of OVLT^Glut^-PVN/RVLM neurons may partly be due to excessive aldosterone. Seven days of spironolactone treatment eliminated the increased AP firing rate caused by current injection in OVLT^Glut^-PVN/RVLM neurons of TASK^−/−^ mice, even though RMP stayed depolarized. Moreover, chronic aldosterone-treated TASK^+/+^ mice showed similar electrophysiological properties in these neurons to TASK^−/−^ mice, despite less membrane depolarization (∼3.2 mV *vs.* ∼6.6 mV in TASK^−/−^ mice). The absence of TASK-1 and TASK-3 channels might also be significant, considering their role in maintaining RMP^64^ and high expression in OVLT, and spironolactone didn’t fully reverse the depolarized RMP in TASK^−/−^ mice. The aldosterone-treated TASK^+/+^ mice exhibited heightened brain salt sensitivity, showing a greater BP response to intracerebroventricular hypertonic solution infusion than controls, which was eliminated by OVLT^Glut^-PVN/RVLM neuron lesions. Notably, neither aldosterone alone nor 1% NaCl alone increased BP in mice, yet their combination effectively raised BP. Immunofluorescence results also confirmed that only the combination effectively activated OVLT^Glut^-PVN/RVLM neurons. Overall, these findings back our hypothesis that aldosterone enhances central sodium sensitivity, through increased excitability of OVLT^Glut^-PVN/RVLM neurons, thereby synergizing with high sodium to elevate BP.

The molecular underpinnings of the observed hyperexcitability in OVLT^Glut^ PVN/RVLM neurons are complex and multifactorial, with aldosterone-induced changes in ion channel expression likely being an important factor. TRPV4 (Trpv4) responds to hypotonic solution and plays a role in thirst and central osmoregulation^65^. Nicotinic acetylcholine receptors (Chrna6, Chrna9) influence RMP control, synaptic transmission modulation, and fast excitatory transmission^66^. Kv12.2 (Kcnh3), a voltage-gated potassium channel, is crucial for regulating neuronal excitability, with its deletion leading to neuronal hyperexcitability^67^. Kir2.3 (Kcnj4) and Kir7.1 (Kcnj13) are inward rectifier potassium channel family important for maintaining RMP and neuronal excitability^68^. Nav1.8 (Scn10a) amplifies neuronal excitability in DRG neurons, especially near the AP threshold, and its increased expression can lead to hyperexcitability and higher firing probability^69^. Therefore, aberrant expression of these genes should be considered the most likely candidate in explaining the increased excitability of OVLT^Glut^-PVN/RVLM neurons in TASK^−/−^ mice. Both TASK-2 (Kcnk7) and ENaC (Scnn1a) are involved in sodium transport. TASK-2 contributes to sodium and water reabsorption by alkalinizing the basolateral membrane, which facilitates sodium reabsorption^70^. ENaC also serves as a taste receptor for sodium^71^ and playing a role in central sodium sensing^72^. In the kidney, aldosterone regulates sodium reabsorption by modulating ENaC. The altered expression of Scnn1a, Scn10a, Chrna6, and Kcnh3 in the OVLT of TASK^−/−^ mice is at least partly attributed to excess aldosterone, as it can be reversed by spironolactone treatment. Moreover, chronic aldosterone treatment in TASK^+/+^ mice induces similar changes. Especially the upregulation ENaC channel mRNA expression may play a role in the central sodium sensing mechanism. Evidence shows intracerebroventricular infusion of ENaC-like channel blockers (amiloride, benzamil) reduces hypertension^73,74^. It is speculated that ENaC-like channels may be in ventricular lining epithelial cells, transporting Na^+^ from cerebrospinal fluid to brain to modulate osmosensory mechanisms^4^. However, the upregulation of Scnn1a (encoding α subunit of ENaC) may not be through the classic aldosterone/MR or GPR30 pathways^75^, given the absence of Hsd11b2 and Gper1, though MR is present in OVLT cells (Fig. S7). Studies indicate MR activation has roles independent of aldosterone levels^76^. Our RNA sequencing and qPCR validations were conducted on the entire OVLT tissue, without verifying whether these dysregulated genes exhibit similar patterns in OVLT^Glut^-PVN/RVLM neurons. Further studies are needed to clarify how aldosterone regulates these gene expressions and their impact on OVLT^Glut^-PVN/RVLM neuron electrophysiology.

### Perspectives

This study identifies a critical role for OVLT^Glut^ neurons in linking aldosterone excess to salt-sensitive hypertension via their hyperexcitability and projections to PVN/RVLM. The findings suggest that aldosterone-induced dysregulation of ion channels (e.g., ENaC and potassium channels) sensitizes these neurons to sodium, amplifying sympathetic outflow and BP. While these mechanistic insights offer potential therapeutic targets—such as modulating OVLT^Glut^ -PVN/RVLM circuitry—key questions remain. Future work should clarify whether adrenocortical-specific TASK-1 and TASK-3 channels deletion recapitulates these effects and explore translational relevance in human hypertension, particularly in primary aldosteronism or salt-sensitive phenotypes. Additionally, investigating sex differences and the interplay between peripheral and central aldosterone actions could further refine this pathway’s clinical significance.

## Author contributions

L.S. and K.L. contribute equally to this work. L.S. conceived and designed the study and drafted the manuscript. K.L. conducted the patch-clamp experiments, and analyzed data; K.Z., W.H., X.Z., J.C. conducted experiments and acquired data; Y.C. conducted the RNAseq bioinformatics; Y.W. generated mice; S.W. and F.Y. supervised and coordinated the study, and revised the manuscript. All authors read and approved the final manuscript.

## Non-standard Abbreviations and Acronyms

OVLT: organum vasculosum of the lamina terminalis
BP: blood pressure
SFO: subfornical organ
MnPO: median preoptic nucleus
PVN: the paraventricular nucleus
RVLM: rostral ventrolateral medulla
aCSF: artificial cerebrospinal fluid
AP: Action potential
AHP: afterhyperpolarization
HR: heart rate
MAP: mean arterial pressure
MR: mineralocorticoid receptor
AVP: arginine vasopressin
TH: tyrosine hydroxylase
TASK: TWIK-related acid sensitive potassium channel
RMP: resting membrane potential
ENaC: epithelial sodium channel

## Acknowledgements

We are grateful to Douglas Bayliss from the University of Virginia for gifting the TASK^−/−^ mice. We also thank the Core Facilities and Centers, Institute of Medicine and Health for experimental and technical support. Graphical abstract was generated using Biorender (www.biorender.com).

## Data availability

The data underlying this article will be shared on reasonable request to the corresponding author.

## Novelty and Relevance

### What Is New?

- First demonstration that OVLT^Glut^ neuron hyperexcitability directly links aldosterone excess to salt-sensitive hypertension.
- Identifies a specific pattern of aldosterone-induced ion channel dysregulation, involving ENaC and potassium channels, as a key mechanism for sensitizing OVLT^Glut^ neuronal hyperexcitability.
- Establishes the OVLT^Glut^-PVN/RVLM neural circuit as a critical pathway in mineralocorticoid-dependent hypertension.

### What Is Relevant?

- Explains how aldosterone exacerbates hypertension through central nervous system mechanisms.
- Provides new insights into salt-sensitive hypertension, a major clinical challenge.

### Clinical/Pathophysiological Implications?

- Reveals potential new therapeutic targets for primary aldosteronism and related hypertensive disorders.
- Supports development of neuromodulatory approaches for salt-sensitive hypertension.

